# Individual heterogeneity and its importance for metapopulation dynamics

**DOI:** 10.1101/2020.08.11.246629

**Authors:** Stefano Masier, Maxime Dahirel, Frederik Mortier, Dries Bonte

## Abstract

Landscape connectedness shapes the exchange of individuals among patches, and hence metapopulation connectivity and dynamics. Connectedness, and its resulting effects on connectivity are therefore rightfully central in conservation biology. However, besides determining demographic fluxes of individuals between patches, connectedness also generates phenotypic sorting and thus impacts local and regional eco-evolutionary dynamics. Despite the central role of connectedness, its effects on individual phenotypic heterogeneity and spatial organization are so far neglected in theory and applications.

Through experimental metapopulations of *Tetranychus urticae* (two-spotted spider mite) with three levels of landscape connectedness and by regularly removing phenotypic structure in a subset of these populations, we tested how regional and local population dynamics are determined both by network connectedness and phenotypic spatial organization.

We find that the self-organizing phenotypic spatial structure increases local equilibrium population sizes and variability. It in contrast dampens the effects of imposed connectedness differences on population sizes and is therefore anticipated to improve metapopulation persistence. Contrary to theoretical expectations, the most locally connected patches within the network showed an overall reduced local population size, possibly originating from a faster depletion of resources from immigrants or transiting individuals.

This experiment shows how metapopulation dynamics can significantly deviate from theoretical expectations due to individual heterogeneity. This potential rescue effect stemming from phenotypical self-organization in space is a key point to consider for conservation actions, especially based on translocations.

## Introduction

Habitat fragmentation has a deep impact on the population structure and demography of almost every species at any trophic level (Ims and Andreassen 1999; Crooks 2002; Cushman 2006; McGuire et al. 2016). Even without any habitat loss, the physical disruption of a previously continuous habitat into a series of patches separated by a more hostile matrix can result in connectivity loss. Connectivity can be defined as the realized dispersal of individuals and/or propagules among patches of suitable habitat. It therefore depends on one end on the connectedness of the landscape, a measure of the average physical proximity among patches that results from the distance between them (Turner 1989; Cushman, McGarigal, and Neel 2008; Wang, Blanchet, and Koper 2014). Connectivity depends on how connectedness interacts with the dispersal capacities of the species involved (Tischendorf and Fahrig 2000). Barring indirect effects of evolution, loss of connectedness is expected to decrease connectivity, reducing the probability that an empty patch will be colonized, or that patches at the brink of extinction will be rescued; overall, it tends to lower the persistence and stability of a metapopulation (Fuller, Doyle, and Strayer 2015; Thompson, Rayfield, and Gonzalez 2017) and is therefore of prime importance for conservation biology.

Connectedness impacts local and regional fluctuations in population sizes, as well as the synchronicity of these changes within the entire metapopulation (De Roissart, Wang, and Bonte 2015; Wang, Haegeman, and Loreau 2015). Theory predicts that lower effective dispersal between disconnected patches raises both local and regional temporal variabilities in population size thereby increasing local patch extinctions while decreasing rescue from immigration (Abbott 2011; Wang, Haegeman, and Loreau 2015; Zhang, DeAngelis, and Ni 2021). On the other hand, a more connected network will also show a higher level of spatial synchronicity (Gouhier, Guichard, and Gonzalez 2010), which makes the whole system also more vulnerable to global extinction because of declines in rescuing potential when all local populations decrease at the same time (Lloyd and May 1999; Sabelis et al. 2005). This holds especially when local patches are small, as small local populations are vulnerable to demographic stochasticity (Fahrig 2017). Recent theory shows, however, a more complicated picture with local variability and synchronicity in these local population fluctuations to depend on the pace of population growth and recovery after perturbance (Luo et al. 2021). Connectivity can thus be considered a double-edged sword for metapopulation conservation (Hudson and Cattadori 1999), but it is clear that more insights are needed on the conditions causing perils for metapopulation persistence.

All these theoretical developments ignore evolutionary dynamics and the potential for eco-evolutionary feedbacks. Landscape connectedness directly affects the costs of dispersal (Bonte et al. 2012), imposing selection pressures on dispersal and life history. The resulting life history and dispersal evolution will in turn feedback on connectivity and demography (Bonte, Masier, and Mortier 2018; Govaert et al. 2019). The evolutionary feedbacks can occur at both the regional (metapopulation) and local (patch) level. Connectivity is an environmental pressure affecting all individuals in a suite of patches which may impact the demography and therefore competition in single patches. Metapopulation dynamics are consequently expected to be influenced by metapopulation-level selection on dispersal and associated traits that may equally evolve at the local patch level. This can include, but is not limited to, feeding efficiency, offspring investment and energy allocation (Masier and Bonte 2020). Indeed, the position and local connectedness of each patch within the spatial network will determine the number of immigrants (Tischendorf and Fahrig 2000; Fahrig 2017; Bonte and Bafort 2019; Karisto and Kisdi 2019). Patches with higher levels of local connectedness are for instance expected to have an increased immigration, that should theoretically increase population densities (Ives et al. 2004; Masier and Bonte 2020). Alternatively, as less connected patches will receive lower numbers of immigrants, they should become genetically more isolated, kin-structured and inbred.

Connectivity not only affects fluxes of individuals among patches, and therefore local and regional population sizes, it is also expected to determine the phenotypic structure within the metapopulation. Changes in local densities affect competition and therefore directly impact the phenotypic signature of the developing individuals (Lion 2018). This phenotypic plasticity is usually expressed in body size, morphology, physiology, and behavior (Parsons et al. 2011; Koeller, Mohn, and Etter 2000; Kamioka and Iwasa 2017; Le Galliard, Paquet, and Mugabo 2015; Woodworth et al. 2017). Since dispersal is performed by a non-random subset of individuals from the population that share specific life history traits (Bonte et al. 2012; Clobert et al. 2012; Dahirel et al. 2019) and physiology (Goossens et al. 2020), connectivity changes will equally affect the sorting of phenotypes, and therefore phenotypic variation (*kind structure*) among patches. The spatial organization of phenotypes among and within patches then again has the potential to affect population dynamics. More-over, because of intimate links to gene flow and thus spatial genetic structure, connectivity will directly impact interactions among relatives (*kin structure*). In absence of realized dispersal, competitive interactions among kin are elevated (Perrin and Mazalov 2000). Kin competition is known to be an important driver of dispersal (Gandon 1999), thus potentially imposing negative feedbacks on the metapopulation dynamics (Poethke, Pfenning, and Hovestadt 2007). At the same time, local interactions between individuals will impose an internal self-organization, with phenotypes exclusively emerging from local demographic interactions (Lowe, Muhlfeld, and Allendorf 2015). This emergent complexity, resulting from a balance between order (phenotypical sorting) and disorder (stochastic distribution of phenotypes) determines the demographic self-organization of the entire metapopulation (Parrott 2010; Urban et al. 2020).

Individual heterogeneity, phenotypic self-organization, selection and the resulting population dynamics are anticipated to impact metapopulation viability (Thompson, Rayfield, and Gonzalez 2017). We used the two-spotted spider mite *Tetranychus urticae* as a model to demonstrate how the spatial self-organization of phenotypes (i.e., the emerging kin and kind structure) equalizes metapopulation dynamics across connectedness levels (Urban et al. 2020). The prerequisites for such a phenotypic self-organization are present: phenotypic variation in relation to local densities, stage and sex structure (Dahirel et al. 2019; De Roissart, Wang, and Bonte 2015), density and kin-competition related dispersal (Van Petegem et al. 2018; Fronhofer, Poethke, and Dieckmann 2015). Moreover, previous studies showed that this species’ phenotype-dependent dispersal maximizes fitness (Dahirel et al. 2019; Bonte et al. 2014). We earlier demonstrated that changes in connectedness impose genetic and epigenetic changes in dispersal timing, dispersal costs and fecundity, but not dispersal propensity (Masier and Bonte, 2020).

We initialized replicated experimental metapopulations according to three levels of metapopulation-level connectedness and implemented a randomization treatment by reshuffling individuals across the patches in the metapopulation, removing the spatial geno- and phenotypic structure while leaving demographic attributes like densities, stage, sex, and age-structure untouched. This treatment thus removed most of the emerging spatial kin and kind structure (Masier and Bonte, 2020). We quantified and compared average local and metapopulation sizes, as well as the level of demographic fluctuations (local and metapopulation variability) and synchrony in local population size changes between treatments (Wang and Loreau 2014). As variation in local connectedness may also affect local dynamics, we additionally compared responses from patches within the same spatial network that varied in their degree of local connectedness (i.e., core vs edge patches within a single metapopulation). We generated some prior predictions on the role of connectedness versus connectivity on metapopulation demography by means of an individual based model, and we later compared them to our experimental results.

## Materials and methods

### Study species

To perform the experiment, we used the two-spotted spider mite *Tetranychus urticae* Koch 1836 (Acari: Tetranychidae) as a model. *T. urticae* is a parasite on a wide diversity of host plants (Migeon, Nouguier, and Dorkeld 2010), and has short developmental time (7 to 10 days at experimental conditions (Hance and Van Impe 1999)). We used the “LS-VL” lab strain established in October 2000 (Van Leeuwen et al. 2008; Cazaux et al. 2014) as the base population of spider mites. This population has been maintained on bean (*Phaseolus vulgaris* L., variety ‘Prelude’) in climate-controlled rooms and under a constant L:D 16:8 light regime since then. As many experiments showed through the years (e.g., Van Petegem *et al*., 2018), this strain shows a high level standing genetic variation and is therefore of perfect use in experimental evolution (Kawecki et al. 2012).

### Experimental setup

To understand the effect of spatial self-organization and spatial connectedness on ecological dynamics, we studied spider mites in 24 independent experimental metapopulations varying in connectedness levels and the application or not of a randomization treatment (see below). These metapopulations are the ones studied in (Masier and Bonte, 2020) for phenotypic evolution; we present here a summary of the experimental setup. Further practical details about the setup and maintenance of the experimental metapopulations are provided in (Masier and Bonte, 2020) and as supplementary material (Supplementary Material S01).

Each experimental metapopulation consisted of a 3-by-3 square grid of nine patches, with connections between patches following Moore neighborhood rules (each patch connected to its “horizontal”, “vertical” and diagonal neighbors). Patches were all the same size (5×5 cm^2^) and were each cut from one of the two main leaves of a two-weeks-old bean plant. Patches were connected using Parafilm bridges; each bridge was about 0.5 cm wide, and the length of the bridges defined the “metapopulation-level connectedness” treatment: 8 metapopulations were closely connected (4cm long bridges), 8 averagely connected (8cm) and 8 loosely connected (16cm). Lengths in the previous sentence refer to all horizontal and vertical bridges; diagonal bridges were ≈ 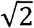 times longer. Experimental trials showed that mortality during movement increased with bridge length (Supplementary Material S02).

Within each replicate metapopulation, patches could be distinguished by their level of “local connectedness”: central patches were connected to all other patches (8 links), while side and corner patches were less connected (5 and 3 links, respectively, (Masier and Bonte, 2020)).

Every week, we counted and collected all adult females using a thin paintbrush, put them on fresh bean leaves to reduce stress as much as possible, after which they were collectively put back in the metapopulation. In the standard treatment used as a control for the reshuffling treatment (see lower), they were put back on the same leaf as the one they were collected from. To tease out the consequences of phenotypic and genotypic heterogeneity (kind and kin structure), we subjected three replicates out of eight in each metapopulation connectedness level to a randomization treatment. In these, collected females were then shuffled and moved back on a random patch of the same metapopulation. Randomization was done so the number of adult females per patch remained unchanged pre- and post-manipulation. As a result, both the absolute population numbers and the stage/sex structure of each patch remained unaltered by randomization itself. The treatment therefore specifically destroys, or at least reduces, kin and kind structure (genetic relatedness and phenotypic similarity; (Van Petegem et al. 2018)) as well as any local adaptation caused i.e. by local density fluctuations, while maintaining the direct ecological effect of connectedness on population densities themselves.

We decided to reshuffle only three out of eight replicates to ensure feasibility over the course of the full experiment (10 months) while optimizing statistical power for both these demographic and earlier evolutionary analyses (Masier and Bonte, 2020).

Each patch was initialized at the start of the experiment using 5 adult females and 2 adult males taken randomly from our stock population (63 individuals per metapopulation, 1512 individuals overall). After a 15-days (≈1.5 generations) “burn-in” period during which no count or randomization was performed to allow (meta-)populations to reach an equilibrium (which was reached out in 21 out of 24 replicates, Supplementary Material S03), the metapopulations were monitored for 23 weeks (≈18 generations at the described conditions). This resulted in a total of 4941 abundance records at the patch level (9 patches × 24 replicates × 23 weeks, minus three replicates missing one week each due to data transcription errors).

### Individual-based model

To understand the importance of connectedness for the measured metapopulation metric, we developed an individual-based version of a metapopulation model that resembled our experimental systems (3×3 patch networks). The model did not incorporate evolutionary dynamics or context (e.g. density-)dependent dispersal. Changes in connectedness were implemented by simulating higher dispersal mortality (as from the experimental result presented in material [Supplementary Material S02]). An extensive model description is provided in the supplementary material (Supplementary Material S04). We present results from this simulation model prior to those observed in the effective experiment.

### Metapopulation dynamics: statistical analysis

Following Wang and Loreau (2014), we can describe the temporal dynamics of each metapopulation by the patch-level average population sizes and a series of four metrics summarizing variability at multiple scales (Table 1). To calculate these metrics and compare them across treatments, one needs to gather estimates of mean population sizes for each patch and temporal covariances between patches for each metapopulation, as well as information about their uncertainties. Wang and Loreau (2014) do not provide ready-made methods to estimate these uncertainties; however, one can readily obtain both point estimates and uncertainties for each of these metrics using the output from a multivariate mixed/multilevel model where each patch is its separate response and patch responses are correlated (Supplementary Material S05).

**Table 1:** Summary of the key variables estimated from the multivariate Poisson model of local population sizes, based on Wang and Loreau (2014) (See Supplementary material S05 for how each metric relates to combination of fixed and random effects).

| Description | Notation |
| --- | --- |
| Temporal variance of local population size in patch $i$ <sup>a</sup> | $w_{ii}$ |
| Temporal mean local population size of patch $i$ <sup>a,b</sup> | $\mu_i$ |
| Temporal (co)variance between patches $i$ and $j$ from the same metapopulation <sup>a</sup> | $w_{ij}$ |
| Temporal mean metapopulation size <sup>c</sup> | $\mu_M = \sum_i \mu_i / m$, where $m$ is the number of patches |
| Temporal variability at the local patch scale $\alpha$: weighted average of patch-level coefficients of temporal variation | $\alpha = \left( \frac{\sum_i \sqrt{w_{ii}}}{\mu_M} \right)^2$ |
| Temporal variability at the metapopulation scale $\gamma$ | $\gamma = \left( \frac{\sum_{i,j} \sqrt{w_{ij}}}{\mu_M} \right)^2$ |
| Additive spatial variability $\beta_2$ | $\beta_2 = \alpha - \gamma$ |
| Spatial synchrony $\varphi = 1/\beta_1$ <sup>d</sup> | $\varphi = \gamma/\alpha$ |

<sup>a</sup> see Supplementary Material S05 and de Villemereuil et al. 2016 for how to obtain this parameter on the data scale from latent-scale mixed model coefficients
<sup>b</sup> It is important to note that in a Poisson model with observation-level random effects, differences in patch-level intercepts reflect differences in (log-transformed) medians; if present, as in here, differences in within-patch variances should be accounted for to estimate differences in means (see de Villemereuil et al. 2016)
<sup>c</sup> Can also be estimated from a model with metapopulation size as a response (see Supplementary Material S07 for a comparison)
<sup>d</sup> Spatial synchrony among patches from the same populations; connects local temporal variability and additive spatial variability to metapopulation temporal variability; $\alpha \times \varphi = \gamma$ , $\alpha \times (1 - \varphi) = \beta_2$ .

Using numbers of adult females per local patch as our metric, we analyzed mite abundance data by fitting a Poisson multivariate multilevel model (Supplementary Material S05). We fitted this model in a Bayesian framework using the Stan language (Carpenter et al. 2017). The use of a MCMC-Bayesian framework is especially valuable here as, among other things, posterior samples greatly facilitate error propagation when combining multiple model parameters to estimate the metrics (described in Table 1). We wrote this model and analyzed it using R, version 4.0.5 (Team R Core 2021) and the *brms* package version 2.14.4 (Bürkner 2018) as frontends. For simplicity, we only present a conceptual summary below: a complete and formal description, including rationale for prior choices and how the metrics described in Table 1 can be derived from model parameters (fixed effects, random effects and (co)variances), is included in the archived code (see “Data and code availability”) and in supplementary material (Supplementary Material S05).

This model included fixed effects of metapopulation-level connectedness, local connectedness, randomization, and the two randomization × connectedness interactions, as well as random effects of metapopulation and patch identity (nested in metapopulation). We converted fixed effects to centered dummy variables to facilitate the interpretation of interaction coefficients (Schielzeth 2010) and the setting of priors. The model also included patch-specific observation-level random effects (Harrison 2014), to account for within-patch temporal variability. These random effects were correlated between patches, to allow for temporal covariance among patches (the *w*_*ij*_ in Table 1). Regarding this within-patch (co)variation, it is important to note that because landscapes can be freely rotated, *x,y* encodings for patch coordinates are somewhat arbitrary and only valid within a given replicate metapopulation. This has important consequences for model specification: indeed, the *x* = 1, *y* = 1 corner patches in any two metapopulations do not intrinsically “match” each other more than, say, the *x* = 1, *y* = 1 corner patch in one landscape and the *x* = 1, *y* = 3 corner patch in another. This prevents explicit in-model partitioning of covariance matrices into mean treatment-level and residual components. As a result, we fitted one within-patch covariance matrix per replicate, and posterior averaging among replicates from a same treatment was only done once all within-replicate operations were complete.

In a multilevel/mixed Poisson model as here, mean population sizes depend on both latent intercepts and within-patch variability (de Villemereuil et al. 2016). Importantly, if the latter random effect variance is allowed to vary between treatments, as is the case here, then we need to account for it to accurately predict average differences due to connectedness and/or reshuffling, as differences based on fixed effects only may not show the whole picture (Table 1). We therefore used formulas in (de Villemereuil et al. 2016) to estimate posterior means on the observed scale from latent-scale intercepts and covariance matrices, and compare them between treatments. We additionally used methods in (de Villemereuil et al. 2016) to convert posterior temporal covariance matrices from our model, which are on the latent log scale, to the observed data scale, which is the scale relevant for the variability metrics (Table 1). We compared mean local population sizes between replicates as a function of connectedness (local and metapopulation-level) and randomization treatment, and connectedness × randomization interaction. For the variability metrics (α, β_2_, ⍰, γ), due to a reduced number of experimental points at the metapopulation scale, we focused solely on the additive effects of connectedness and randomization.

In a secondary model, we analyzed mean metapopulation-level abundances between treatments, i.e., the response variable was here the sum of adult females in one replicate. Because there was no internal covariation to worry about at that scale, we used a negative binomial model, in which both the mean abundance and the within-metapopulation temporal variability (through the shape parameter) depended on fixed effects of landscape connectedness, randomization and their interaction, and random effects of metapopulation identity. We compared posterior of treatment effects on one side, and mean replicate-level metapopulation size on the other side, between this model and the main one (in the latter multiplying local predictions by the number of patches) to get metapopulation sizes.

Priors choices for both models combined general-purpose “weakly informative” priors based on (McElreath 2020), with more informative priors for parameters where pre-existing knowledge was strong (based on De Roissart, Wang, and Bonte 2015), or where constraints on model parameters were needed (Gelman et al. 2020). We based the more informative priors (fixed effect intercepts and random effect variance) on the empirical distribution of population densities in (De Roissart, Wang, and Bonte 2015). For each model, we ran four chains for 4000 iterations, with 2000 iterations post-warmup per chain; this ensured both convergence (based on graphical checks and the modified 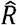 statistic (Vehtari et al. 2020)), and that bulk and tail effective sample sizes were satisfactory for all parameters ((Vehtari et al. 2020); at least 400, and at least 1000 for most parameters). We relied mainly on the *tidybayes, bayesplot, patchwork* packages as well as the *tidyverse* suite of packages for data handling, model post-processing and plotting (Gabry et al. 2019; Pedersen 2019; Wickham et al. 2019; Kay and Mastny 2020). All posterior summaries given in text and figures are given as means [95% Higher posterior Density Intervals (HDIs)].

## Results

### 1. local and regional population sizes

From the simulation model, we expect population sizes to consistently decrease with a decreased regional (metapopulation) connectedness (Supplementary Material S04). Patches with higher local connectedness (central location) harbored consistently larger populations than less connected patches (corner location) (Supplementary Material S04).

Randomization, local and metapopulation connectedness impacted the populations sizes in the experimental metapopulations. The directions of effect were however different. For the additive effects, less connected metapopulations had higher mean population sizes (Fig. 1; predicted ratio 16 cm / 4cm = 117.2% [102.9%, 132.9%]). While evidence is weaker (95% interval overlaps 0), the direction of response is the same in response to local connectedness (Fig. 1, predicted ratio 3 links / 8 links = 113.5% [98.3%, 127.2%]). Local populations were on average smaller in reshuffled metapopulations compared to controls (Fig. 1, predicted ratio reshuffled/control = 87.0% [78.0%, 95.9%]).

**Figure 1.**
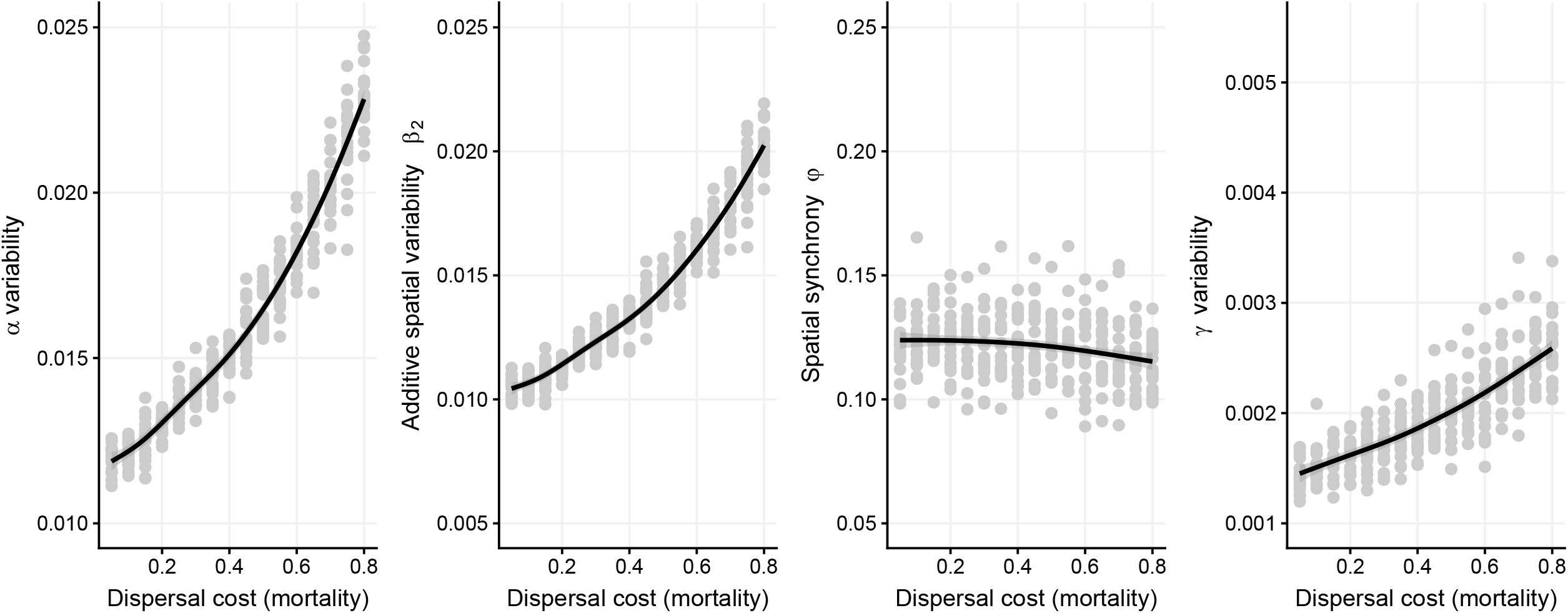
Posterior additive effects of treatments on mean local population size. For each treatment variable, posteriors (with mean and 95% CI in black) are displayed after averaging over the other two. Grey dots are empirical patch-level means, transparency of the dots is proportional to relative variability, with more precise estimates being more opaque (using the inverse of the mean-scaled variance as a measure of precision; see e.g. (Dochtermann and Royauté 2019)).

Looking separately at the randomized versus non-randomized metapopulations, possible interactions became clear (Fig 2). The effect of metapopulation-level connectedness on population size was mostly apparent in the randomized metapopulations (predicted ratio 16cm/4 cm = 131.8% [106.4%, 157.8%]) but strongly reduced when looking only at control replicates (predicted ratio 16cm/4 cm = 110.1% [93.1%, 127.4%]). Conversely, the effect of the randomization treatment on population sizes was mostly detected at high levels of connectedness, but not at the lowest (randomization/ control ratios in 4 cm landscapes: 84.7% [70.3%, 99.5%], in 8 cm landscapes: 74.6% [61.5%, 88.5%], in 16 cm landscapes: 101.1% [84.4%, 119.9%]). For a more detailed comparison between posterior distributions, please refer to the Supplementary material (Supplementary Material S06)

**Figure 2.**
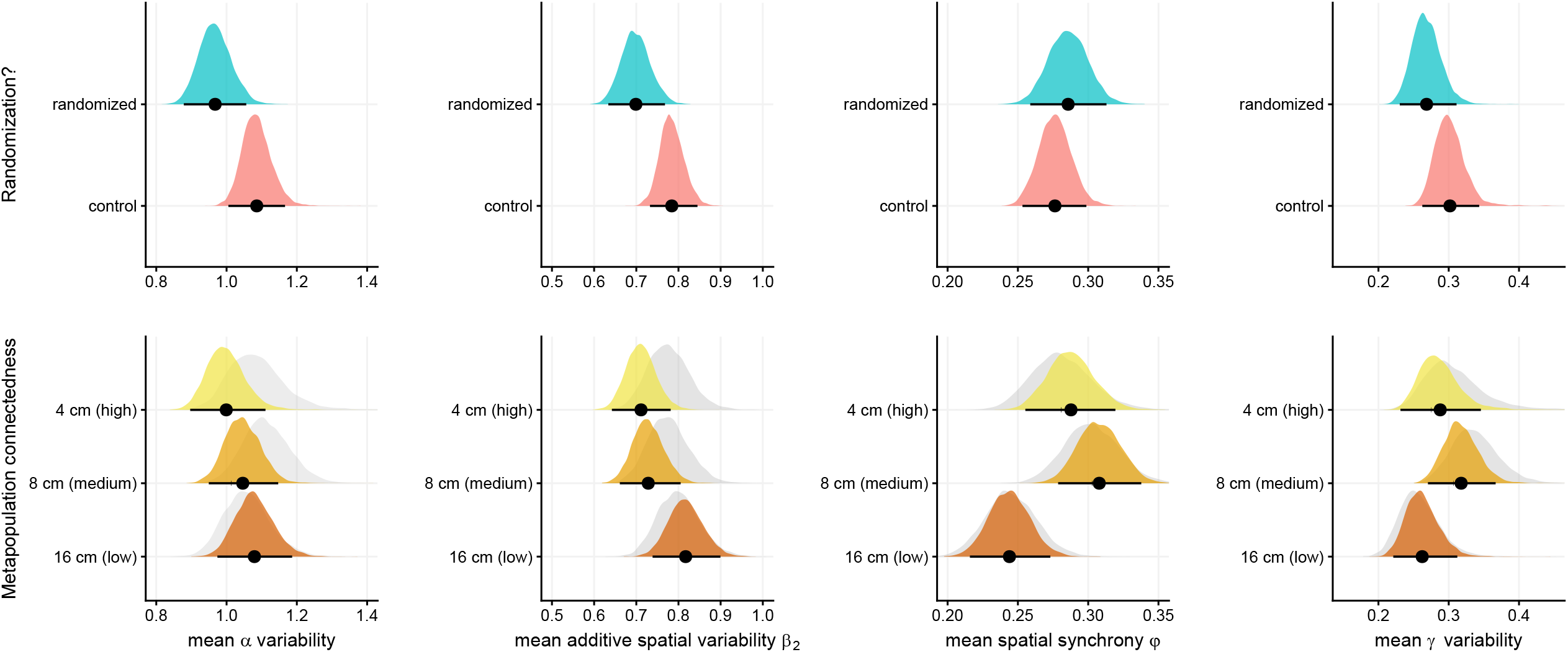
Posterior effect of metapopulation connectedness on mean local population size as a function of randomization. For each connectedness level, posteriors (with mean and 95% CI in black) are displayed after averaging over the other one. Grey dots are empirical patch-level means, transparency of the dots is proportional to relative variability, with more precise estimates being more opaque (using the inverse of the mean-scaled variance as a measure of precision).

Finally, the analysis of mean metapopulation sizes in a separate model shows no clear effects of any treatment (Supplementary Material S07). However, although all effect size posteriors overlap 0, possibly due to the much lower sample size at that scale, the directions of effect are nonetheless the same as those implied by the patch-level model (Supplementary Material S07), and mean replicate-level predictions are congruent with those that would be obtained by summing local estimates from the patch-level model (Supplementary Material S07).

### 2. Metapopulation variability

Our simulations showed that both local (α) and regional (γ) variability are expected to increase as a direct result of higher dispersal mortality under connectedness loss (Fig.3). Synchrony (1), by contrast, remained unaffected (Fig.3).

**Figure 3.**
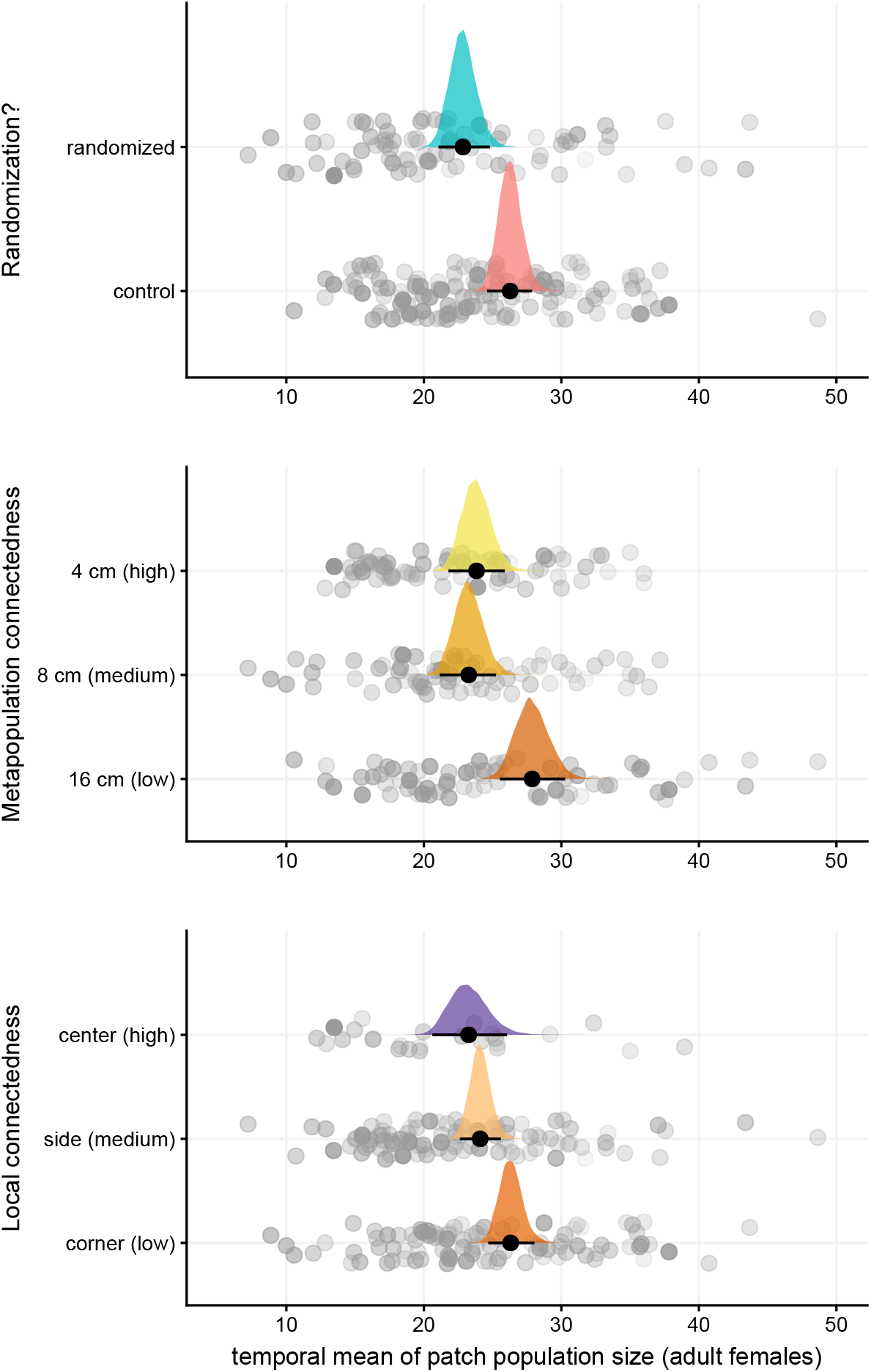
Landscape variability metrics calculated from our simulation model (no evolution included, dispersal mortality used as a proxy for connectedness): variability is expected to increase with lower connectedness, both at local (first panel) and metapopulation level (fourth panel); spatial variability is expected to increase similarly (second panel). Conversely, spatial synchrony is expected to be largely unaffected (third panel).

In our experiments, reducing metapopulation-level connectedness reduced spatial synchrony (predicted ratio 16 cm/4cm = 85.1% [71.7%, 99.0%]) and increased additive spatial variability (115.2% [99.7%, 131.4%]), but had no effect on α or γ variability (Fig. 4). By contrast, randomization had a possible dampening effect on both local (α) variability (predicted ratio reshuffled/control= 89.2% [78.3%, 99.8%]), and additive spatial variation (89.3% [79.0%, 100.3%]), but no detectable effect on synchrony or metapopulation-scale (γ) variability (Fig. 4).

**Figure 4.**
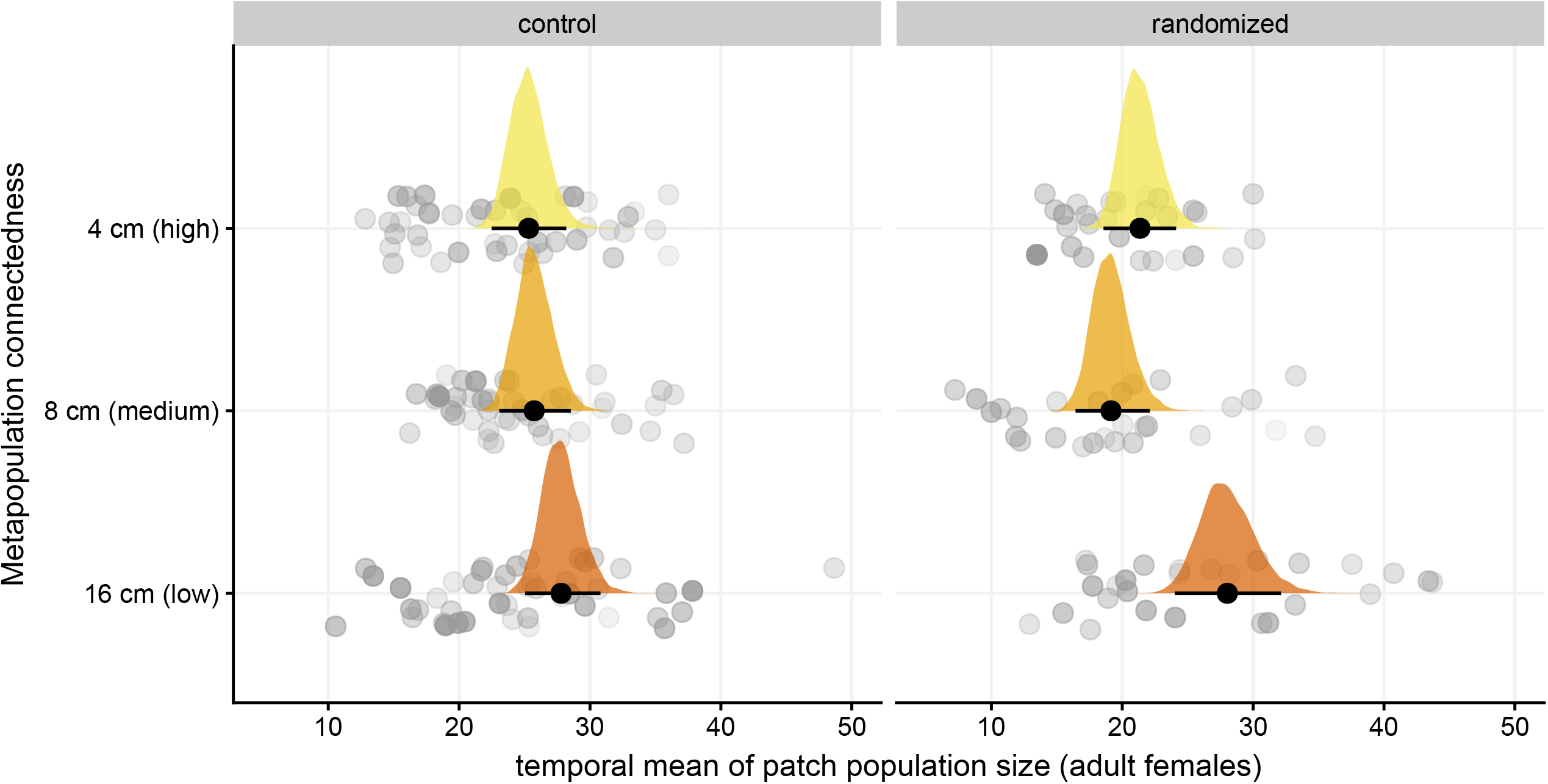
Effect of landscape-level variables of measures of landscape variability based on (Wang and Loreau, 2014). For each treatment variable, posteriors (with mean and 95% CI in black) are displayed after averaging over the other two. Grey “shadows” in the panels showing the effect of connectedness denote the posteriors that would be obtained using only control, non-randomized replicates.

## Discussion

Metapopulation dynamics, as summarized by average population size, population size variability, spatial unevenness and spatial synchrony are known to be impacted by the level of connectedness (Wang and Loreau 2014; De Roissart, Wang, and Bonte 2015; Wang, Haegeman, and Loreau 2015), dispersal (Jacob et al. 2019; Mortier et al. 2019) and the interaction of dispersal with local environmental and demographic variation (Fronhofer et al. 2018). We earlier quantified trait evolution in response to changes in connectedness (Masier and Bonte, 2020) and now show that phenotypic sorting and spatial organization can counter the expected decrease of population sizes loss, thereby potentially safeguarding metapopulation stability. Dispersal in general is phenotype-dependent (Clobert et al. 2009; Bonte and Dahirel 2017), leading to the sorting of phenotypes in metapopulations (Bonte et al. 2014; Dahirel et al. 2019). We provide experimental evidence that such a phenotypic self-organization is an additional and powerful driver of metapopulation dynamics. The exact underlying suites of trait changes that drive this pattern remain, however, unknown.

Theory predicts that lower connectedness and dispersal intensify local and regional variabilities in population sizes (Singh et al. 2004) and increase spatial asynchrony within the metapopulation (Wang, Haegeman, and Loreau 2015). The former is also supported by the simple individual-based model inspired by our experimental setup (Fig.3). We did, however, not observe such a response in our experiments, and if anything, the opposite was found with higher levels of synchrony emerging in the least connected metapopulations (Fig. 4).

Our experimental results can be reconciled with the abundant theory on the synchronizing effects of dispersal in several ways. First, connectedness loss can in some cases drive the evolution of higher dispersal (Cote et al. 2017) which then compensates the desynchronizing effects of connectedness loss. Dispersal is known to often be positive-density dependent, both in general (Harman et al. 2020) and in spider mites specifically (Bitume et al. 2013). It is worth remembering that, as local density is not a fixed characteristic of patches, purely density-dependent habitat choice fails to promote local evolutionary differences (Berner and Thibert-Plante 2015). We expect a stronger local density-dependency when dispersal costs are higher (as in our least connected treatments, Supplementary Material S04) (Travis, Murrell, and Dytham 1999; Rodrigues and Johnstone 2014; Govindan et al. 2015). Our results here suggest that because of dispersal variability, landscapes with low connectedness could thus counter-intuitively lead to higher, and more homogenizing movement between patches than expected. Second, if changes in connectedness lead to sufficiently different ecological conditions through feedbacks, then individuals may plastically adjust their responses and eventually show higher dispersal, but also on average higher fecundity and growth rates (Dahirel et al. 2019) that buffer local population fluctuations. Finally, synchrony and variability may simply show opposite patterns at long *versus* short time scales, depending on population growth responses to perturbations (Luo et al. 2021). As such, our findings do match theoretical predictions of dispersal (in our case connectivity) destabilizing populations and desynchronizing population dynamics across patches at short time intervals for populations exhibiting under-compensatory growth (i.e. slow recovery after being perturbed). At longer time intervals, these dynamics shift towards the anticipated synchronizing and stabilizing dynamics as found in our theoretical model. Evidently, such long-term experiments are challenging using model species with generation times of larger magnitudes than protists and other microbial models.

Spatial heterogeneity in demographic rates may alter and even reverse population-level predictions from mean-field theory (Holt 1985; Van Dyken and Zhang 2019; Deangelis et al. 2020). When resource conditions are homogeneous among patches, as in our experiment, such a spatial variation in demography can only result from phenotypic structuring and self-sorting. This spatial heterogeneity can be generated by for instance local differences in competition, or any associated phenotypic and genotypic sorting in the network (Masier and Bonte, 2020). Our randomization treatment, where population densities and stage structure were maintained but not genetic (kin) and phenotypic (kind) structure (Van Petegem et al. 2018) thus provides further insights on the relevance of phenotypic heterogeneity.

Disrupting phenotypic self-organization by randomization seems to have negative effects on local populations, based on population sizes (Fig. 1). A closer look reveals that this effect is actually connectedness-dependent (Fig. 2), or conversely that the effect of connectedness is randomization-dependent: randomization has little to no effect in the least connected metapopulations, while it does decrease population size in the others. Phenotypic self-organization thus increases population size above the baseline expected with global random dispersal when connectedness is high. The balance between the loss of group-living benefits (Oku, Magalhães, and Dicke 2009; Yano 2012; Strodl and Schausberger 2013) and the spread of favorable genotypes provides a tentative explanation. When costs are low (high connectedness), randomization is expected to destroy favorable local structure without clear benefits in terms of costs avoided, leading to reduced population sizes. When connectedness is low however, the balance would be positive or neutral, as lost benefits are compensated by reduced costs of dispersal, randomization acting as a no-costs source of global dispersal.

Besides the earlier discussed effect on population sizes, randomization also decreased local (α) and to a lower extent regional (γ) variability (Fig. 4). This shows that (meta-)population variability shifts in the directions anticipated from ecological theory when spatial individual heterogeneity in metapopulation dynamics is removed (Vindenes, Engen, and Sæther 2008; Gangloff, Sparkman, and Bronikowski 2018; Hamel et al. 2018; Smallegange, Fernandes, and Croll 2018). Disrupting spatial phenotypic heterogeneity through randomization decreases in consequence both population sizes and their variability across the connectedness gradient. Or, put alternatively, the maintenance of this phenotypic organization leads to more (asynchronous) fluctuating metapopulations of larger size, conditions known to rescue metapopulations at intermediate levels (Wang and Loreau 2014; Wang, Haegeman, and Loreau 2015), but that can also potentially lead the metapopulations to be more vulnerable towards local extinctions if said fluctuations become too pronounced or erratic (Abbott 2011; Zhang, DeAngelis, and Ni 2021).

Spatial synchrony (⍰) remains unaffected by the randomization treatment. In conjunction with the effects on spatial variability (additive β), we propose the following explanation for this paradoxical result. Spatial synchrony ⍰ *sensu* Wang & Loreau (2014) represents roughly how the regional variability γ is partitioned between local variability α and inter-patches spatial variability β (see also Table 1). As such, if local variability and spatial variability vary with similar dynamics, the resulting spatial synchrony will remain unchanged. This is visible in of simulation model (fig.3): local variability (α) and spatial variability (β) have an almost-identical shape, resulting in a constant spatial synchrony (⍰). In the same regard, the effect of randomization on spatial variability is similar to the one for local variability (fig.4). To summarize, when phenotypic self-sorting is removed from a (meta-)population, local population sizes decrease, while the spatial synchronicity (⍰) remains unaffected because both local variability (α) and spatial variability (additive β) change in the same direction. Self-sorting thus increases both local variability and spatial variability. They respectively impose a destabilizing and stabilizing effect, with the net balance seeming overall positive (at least in terms of population sizes). Further studies are recommended to confirm or falsify our finding, but also to test the consistency of these dynamics on longer time periods, or under disturbances (Luo et al. 2021).

Our experiments additionally show how differences in connectedness within the spatial network affect local population sizes. As the central patches showed the highest level of connectedness in all metapopulations, increased population sizes due to positive immigration-emigration balances were expected (Masier and Bonte, 2020). Surprisingly however, opposite patterns were found with central patches systematically showing lower densities. This suggests that local demographic changes are not captured by the metapopulation-level rates of dispersal. As we did not record the total number of individuals, but only the adult females, it is possible that juveniles development was slowed down in central patches due to resource limitation, which was possibly caused by a large influx of immigrants that since then died or dispersed again. It is known that *T. urticae* exhibits longer development times when faced with suboptimal food sources, while longevity and egg-laying performances are not affected (Aucejo-Romero, Gómez-Cadenas, and Jacas-Miret 2004). Likely such delays caused a reduced number of adults during the counts of central patches relative to the others. Highly-connected patches are commonly considered as the most resilient in a network precisely due to their high number of connections, and thus the most valuable and cost-effective populations for conservation (Calabrese and Fagan 2004; Gonzalez, Thompson, and Loreau 2017; Liao, Shen, and Liao 2020). Our experiment shows how this is not necessarily the case.

The implemented randomization treatment can be considered as an extreme case of hypermobility within the spatial network. Our experiment thus pinpoints the applied consequences of hypermobility, for instance through human-mediated dispersal (McRae et al. 2012; Bullock et al. 2018; Rudnick et al. 2012). Apart from the potential to impose maladaptation, or to hinder adaptation (Brady, Bolnick, Barrett, et al. 2019; Brady, Bolnick, Angert, et al. 2019), hypermobility can lead to changes in the local and regional population dynamics. While phenotypic variation has been shown to explain patterns in biodiversity (Bolnick et al. 2011; Dahirel et al. 2019), we here demonstrate its importance for metapopulation demography, and in extension metapopulation conservation. Removing spatial phenotypic organization by for instance assisted migration needs to be done with care, as it could have so far unforeseen detrimental effects for the persistence of the metapopulation.

