## Supplementary Material 1-7 for "Individual heterogeneity and its importance for metapopulation dynamics"

#### S01 – Experimental metapopulations – practical details and maintenance during experiments

All beans leaves used in the experiment were sourced from bean plants grown without using any pesticide and under controlled climatic rooms (21 ± 1°C, L:D 16:8).

Each metapopulation was independently mounted on a bed of wet cotton, which was kept soaked with distilled water to provide hydration to the leaves and to create a barrier to prevent the mites from escaping or moving between patches outside the designated routes. To ensure constant hydration, automatic gardening pumps (GARDENA® set 1265-20) were connected to a reserve of distilled water and set to deliver a fixed amount of water through pipes during 1 minute once every 8 hours. The water flow was monitored to ensure that it never flooded the leaves or the bridges, to prevent additional stress and ensure that the connection between patches was always maintained.

Landscapes were contained in top-opened plastic crates whose sides were covered in a thick Vaseline layer, to further ensure that no cross-contamination could happen between different metapopulations. The water pipes were coated in Vaseline as well, and were positioned so that there was as little direct contact with the cotton as possible, to reduce the chance of mites climbing on them. To ensure resources remained fresh throughout the experiment, every leaf in the setup was refreshed with a new one once a week, at the same time counting was done. First, adult females were counted on the leaf under a stereomicroscope; then they were moved to the new leaf squares, following the protocol described in the main text. Finally, the old leaves were laid above the new ones, sustained by short sticks to reduce direct contact (to insure ventilation and prevent the spread of molds and fungal infestations), and left in place for 48 hours to allow males and juveniles to move on the new leaves as well. By keeping the old leaves detached from the cotton, we also induced a quick desiccation and a sudden drop of the food quality, thus stimulating the dispersal of the individuals towards the new leaf. After 48 hours, all the old leaves were removed along with the sticks to prevent mites from spinning webs around them.

#### S02 – Quantification of connectedness-dependent dispersal mortality

We tested how dispersal success and individual survival is affected by the length of a Parafilm bridge in a setup similar to the one described in the main text, to empirically demonstrate that the lengths we chose in the main experiment do apply a significant selective pressure onto dispersers.

We tested 4, 8, 16 and 32 cm-long bridges, with five independent replicates per length. The plastic bridges were mounted on a bed of cotton, kept wet using abundant distilled water, and connected on one end only to a fresh bean (*Phaseolus vulgaris* L. cv. Prélude) leaf cut (2.5 × 1.5 $cm^2$ rectangle) to provide secure shelters and food sources to the mites. At the opposite ends of each bridge, we placed 10 adult females from our LS-VL stock population using a thin pen brush. Female age was not controlled for, to better approximate a real, non-synchronized population. Females were placed on a "waiting area," delimited by a strip of wet paper placed orthogonally onto the bridge approximately 2 cm from the leaf-free end, in order to stop female dispersal until all individuals had been placed. Similar strips were used to lock the leaves in place on the wet cotton and to the bridges, as well as at the open end of the plastic bridge to keep it adherent to the wet cotton. No paper strips were placed along the bridge itself, so dispersing mites were not prevented from falling into wet cotton. All replicates were initialized at the same time, by removing the paper strips that stopped movement; the bridges were then stored into a climatically controlled room for 24 hours ($\simeq$ 25°C, L:D 16:8). We then counted the number of live mites on the bridges themselves and the arrival leaves; every unaccounted individual was considered dead and sunken into the wet cotton. We analyzed the effect of bridge length (as a continuous variable) on the proportion of dead individuals using a binomial generalized linear model with weakly informative priors (Normal$(0, 1.5)$ for the intercept and Normal$(0, 1)$ for the slope)(McElreath 2020).

##### Results

The proportion of dead individuals increased with the length of the plastic bridge the mites were placed upon ($\beta = 0.04$ [0.02, 0.07], **Fig. S02.1**).

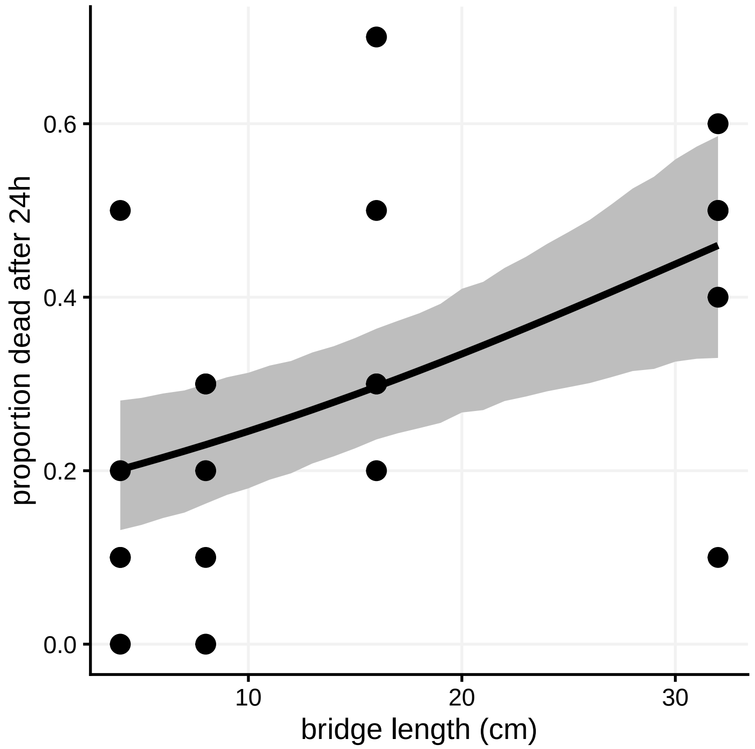


**Figure S02.1** – Mortality rate as a function of bridge length during preliminary trials. Both observed values (dots) and posterior means with 95% credible band are plotted.

#### S03 – Stationarity in experimental metapopulations

A key prerequisite to the use of the variability metrics in Wang and Loreau (2014) is stationarity. To check for stationarity in our metapopulations, we fit a negative binomial generalized linear model including metapopulation-specific intercepts and effects of time (priors are the same as for the negative binomial model described **Supplementary Material S05**). In the large majority of the metapopulations (21 out of 24), we find no evidence of deviations from stationarity (no temporal trend) during the experimental run (**Figure S03.1**).

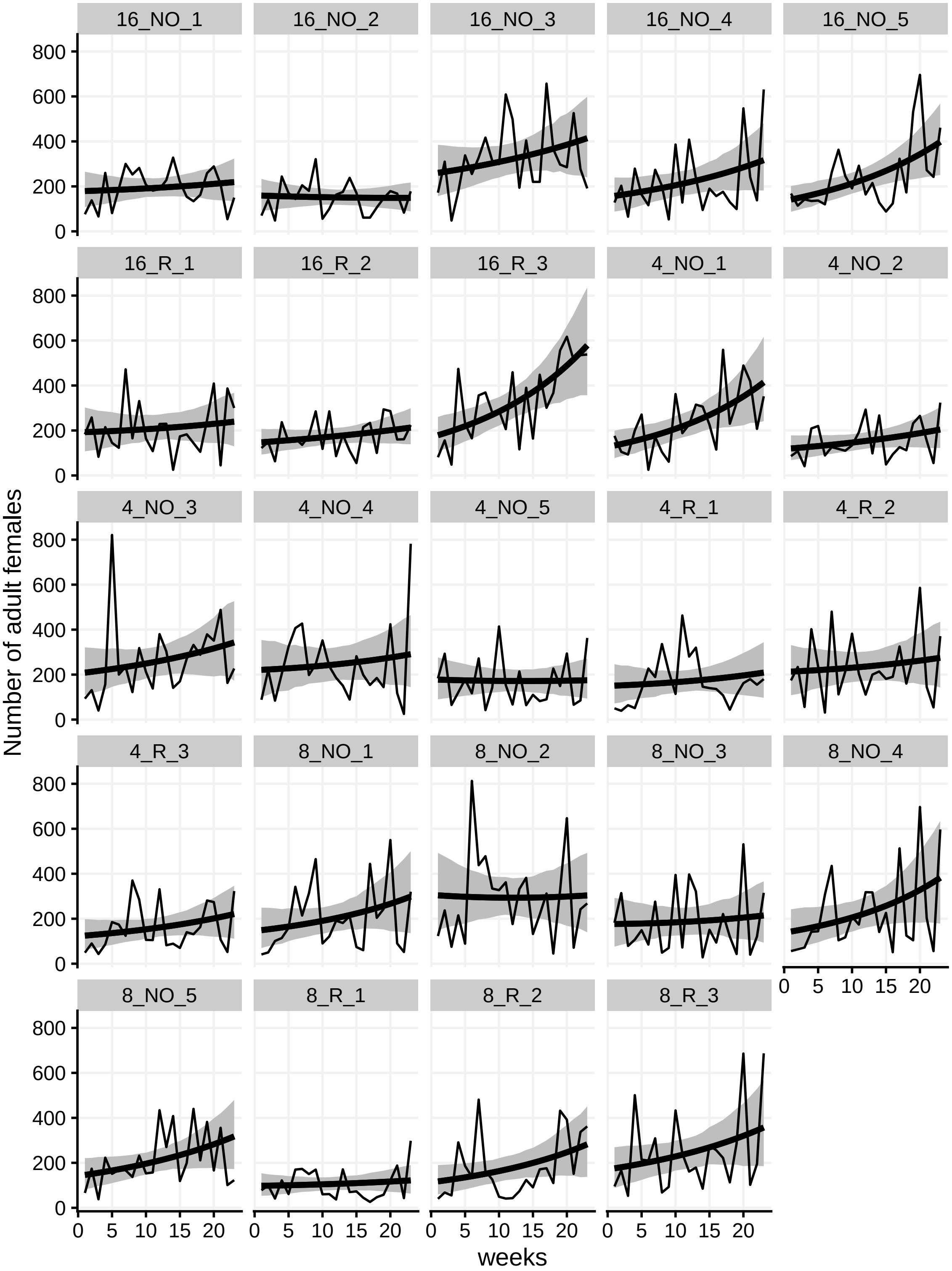


**Figure S03.1** – Metapopulation sizes (number of adult females in all 9 patches of a metapopulation) as a function of time. Both observed values (thin lines) and the posterior means with 95% credible band are plotted. Each subplot represents a different replicate metapopulation.

### S04 – Simulation model description and interpretation

We simulate the direct effect of connectedness on metapopulation variabilities to contrast with the observed dynamics in the experiment, in order to estimate direct ecological impact of the landscape composition in our experiment. We simulated metapopulations resembling our experimental systems, 3×3 connected patch network, in Python (3.8.5). We constructed an individual-based version of the classic metapopulation model in discrete generations. Individuals do not differ in traits within a simulation and we only simulate dispersal and reproduction. Evolution within a simulation is excluded. We analyzed different levels of dispersal mortality to simulate higher dispersal costs and analyze the direct effect of a lower connectedness in our metapopulations (see **Fig. S02.1**). We decided to use mortality as a super-parameter that encompasses both the direct (i.e. death during dispersal) and the deferred (i.e. reduced reproductive success due to energy expenditure) costs of the dispersal event (Bach et al. 2006).

#### ODD description

We describe here the individual-based model used in the main text, using the ODD (Overview, Design, Details) protocol (Grimm et al. 2010).

##### Purpose

The model simulates identical individuals moving and reproducing in a network of nine habitat patches in order to generate metapopulation dynamics, and obtain information about $\alpha$-, $\beta$- and $\gamma$-variability following Wang and Loreau (2014). We simulate what metapopulation dynamics would emerge if a population of a certain phenotype moves as a metapopulation through a certain patchy landscape. Most notably, we simulate what dynamics would emerge independent of individual heterogeneity, density-dependent dispersal, or evolution. The network was simulated to resemble the structure of the metapopulation mesocosms in the main text. We then analyze what the effect of dispersal mortality (as a characteristic of the landscape) is on metapopulation dynamics.

##### State variables and scale

The model simulates individuals as the smallest unit. Individuals only vary in their location in the landscape, which can change to represent movement. They inhabit metapopulations, which is another modeled unit that represent both the landscape and characteristics of the population living in it. All metapopulation characteristics are fixed within a simulation. Dispersal mortality vary across simulations in the range [0.05, 0.80]. Each independent landscape contains nine patches arranged in a 3×3 square and connected following Moore neighborhood rules, with each habitat patch being of undefined size. Landscapes were run for 500 time steps with each time step being a discrete generation.

##### Process, overview and scheduling

Each simulation, after initialization, runs through 500 time steps. Each time step is a discrete generation where individuals, in a random order, each run through their life cycle. A life cycle in the model starts with reproduction, which includes population regulation based on the local population size. After that, the offspring disperses according to a certain propensity. Dispersal includes dispersal mortality and updates that offspring's location ($x$- and $y$-coordinates) if it is successful. This means that dispersal happens before reproduction in the life of an individual (even if in the model's definition of a life cycle, dispersal of the next generation is already modeled while the current one's runs). When all individuals went through their life cycle and died, local population sizes of the offspring are recorded for that generation.

#### Design concepts

**Emergence**: the number and distribution of individuals emerges from movement according along the network and population regulation on reproduction and therefore local population sizes.

**Interaction**: Individuals only interact with one another when in the same habitat patch by competing for local resources. This is included as a population regulating reaction norm that determines reproduction.

**Stochasticity**: Stochasticity plays at many points in the model. At initialization, individuals are randomly placed in one of the habitat patches to generate a random starting distribution. Each individual disperses with a probability according to the dispersal propensity. If it does, it dies with a probability according to the dispersal mortality. If it survives, it disperses to one of the possible destination according to the network with an equal probability for each destination. Each of these are individual events that collectively should result in a certain random portion of the individuals doing one option. An individual's realized number of offspring is drawn from a Poisson distribution with the expected number of offspring as its mean. This simulates a certain variability in fitness and conveniently returns an integer from a decimal expected number of offspring. The sequence of individuals that run through their life cycle is randomly chosen each generation.

**Observation**: Population sizes for each habitat patch is recorded at the end of each generation. From that we calculate metapopulation size for each generation, average metapopulation size, average local population size for each location in the network, $\alpha$-, $\beta$- and $\gamma$-variability, as well as spatial synchrony $\varphi$.

**Scheduling**: Time is modeled in discrete time steps representing discrete generations. During each generation every individual runs through its life cycle before the next individual starts its life cycle. Earlier individuals do not change the environment in a way that would affect later individuals in the sequence.

#### Initialization

At the start of a simulation, a metapopulation is initialized with parameters that will stay fixed for the duration of that simulation. The only parameter that varies among simulations is the dispersal mortality. Furthermore, the model initializes a population with half the metapopulation carrying capacity. This is half of nine times the carrying capacity of a patch or 450 ($K = 100$, **Table S05.1**) individuals that are randomly allocated to one of the nine patches.

#### Input

Each simulation receives a dispersal mortality ($m$) parameter that ranges from 0.05 to 0.8 in steps of 0.05 (**Table S05.1**). We choose not to simulate dispersal mortalities above 0.8 since that would simulate something closer to an unconnected network with a high mortality that are prone to population crashes. We replicate simulations of every input value twenty times.

#### Submodels

**Reproduction** – Each individual $i$ reproduces resulting in a number of offspring ($f_i$) which is drawn from a Poisson distribution:

$$f_i \sim \text{Poisson}(\mu_i).$$

$\mu$ is the expected number of offspring that is affected by the current number of adults, already produced offspring of that generation not included, according to Hassell's population model:

$$\mu_i = r\left(\frac{1+N_i(r-1)}{a}\right)^b,$$

with $r$ the optimal population growth parameter, $N_i$ the local population density experienced by individual $i$ (excluding newly generated offspring), $b$ the shape parameter that determines the type of modeled competition and $a$ the population regulation parameter that is related to carrying capacity ($K$):

$$a = \frac{K(r-1)}{r^{1/b}-1}.$$

An individual is born in the same patch as its parent. Parameters are fixed for all runs (**Table S05.1**).

**Dispersal** – Every newly generated offspring disperses with a probability equal to the simulation's dispersal propensity ($d$). If it disperses, it dies and is not added to the population of the next generation with a probability equal to the simulation's dispersal mortality ($m$). If the dispersing individual does survive dispersal, it will change location to one of the possible destination patches that has a direct link with the patch the offspring was born in at random.

**Table S04.1** Model parameters.

| Symbol | Description | Values |
|---|---|---|
| $r$ | Optimal growth rate | 2 |
| $b$ | Hassell shape parameter | 2 |
| $K$ | Local carrying capacity | 100 |
| $d$ | Dispersal propensity | 0.25 |
| $m$ | Dispersal mortality | 0.05, 0.1, 0.15, 0.2, 0.25, 0.3, 0.35, 0.4, 0.45, 0.5, 0.55, 0.6, 0.65, 0.7, 0.75, 0.8 |

#### Results and interpretation

The simulation model showed local ($\alpha$) and regional ($\gamma$) variability are expected to increase as a direct result of higher dispersal mortality under connectedness loss. As a results of spatial variability ($\beta_2$) increasing at the same time, spatial synchrony ($1/\beta$) remained unaffected (main text, **Fig. 3**). This is opposite to the observed metapopulation variabilities in the experiment. There, local ($\alpha$) and regional ($\gamma$) variability showed no effect of the connectedness treatment while synchrony ($1/\beta$) increased with increasing connectedness. Therefore, heterogeneity in the observed dynamics across treatments cannot only be explained by the direct effect of dispersal mortality or similar dispersal costs alone. Other effects such as (dispersal) trait evolution, and/or the consequences of density-dependent dispersal likely affected metapopulation variabilities meaningfully.

Simulated metapopulation size decreases with increasing dispersal mortality under connectedness loss (**Fig. S04.1**, top) while we observed a higher metapopulation size in the least connected metapopulation when randomized and equal metapopulation sizes when not randomized. Simulated metapopulations showed biggest local sizes in the central patch and lowest in the corner patch (**Fig. S04.1**, bottom), which was the reverse of what was observed. Again, other effects such as genetic structure in the metapopulation or a stronger population regulation effect of resource depletion in the experimental setup likely affected this inconsistency.

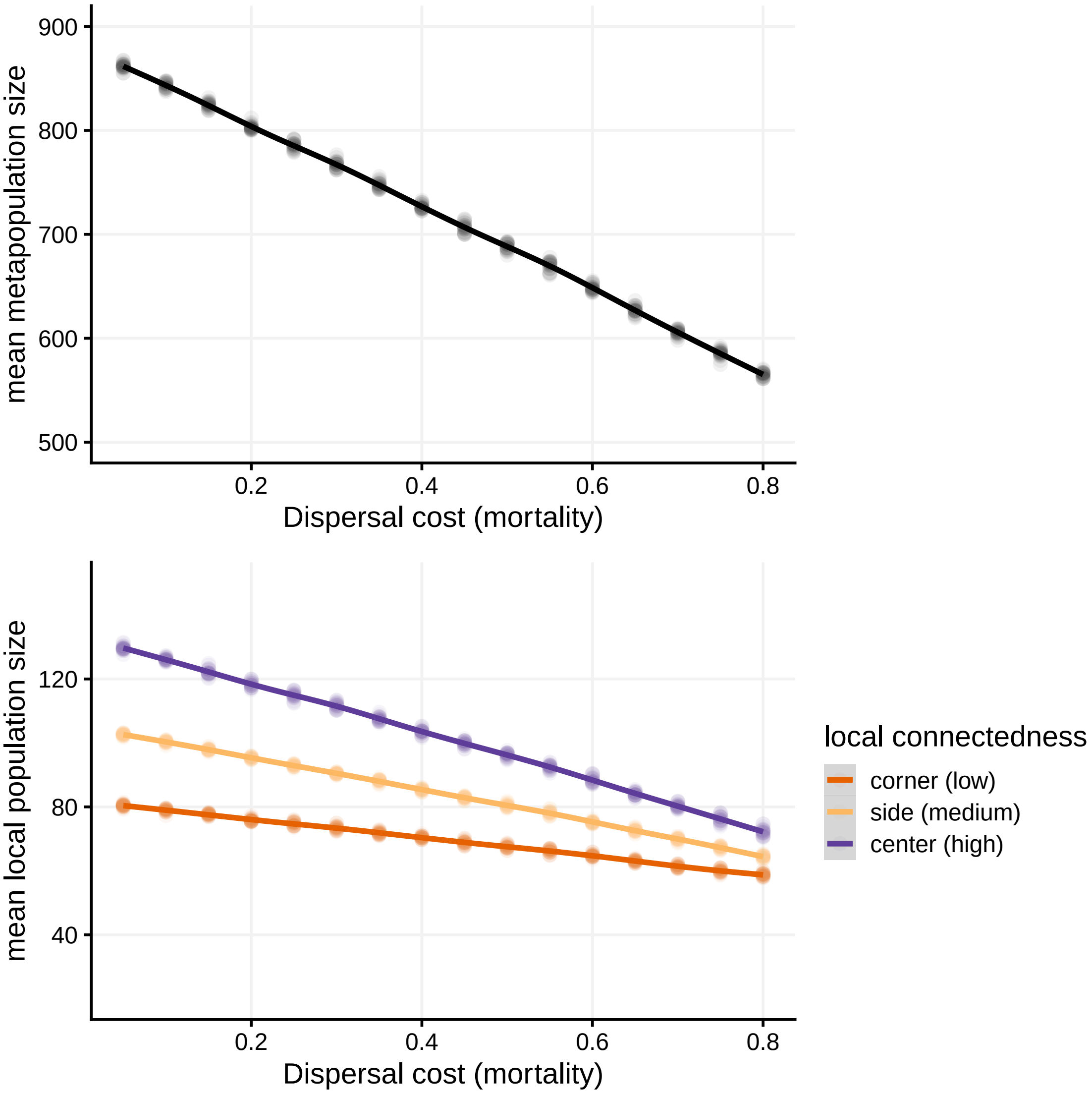


**Figure S04.1** – Effect of dispersal mortality on mean metapopulation and local population sizes, in simulated metapopulations

### S05 – Models description

#### Patch-level model

We fit a generalized linear mixed/multilevel model to the abundance data. This Poisson model (because of count data) includes "random" effects of metapopulation ID (to account for the fact some replicates may have higher/lower average population sizes) and patch nested in ID (because patches may differ beyond the effect of treatment, local connectedness and replicate). Importantly, it also includes a time random effect, to account for temporal patch variance-covariance. You can see below that we are estimating a separate temporal variance-covariance matrix for each replicate $i$. This means that (i) each patch has its own temporal variance, and patches from the same replicate can be correlated. We average everything downstream of the model as needed, but given the non-linearities everywhere in a GLMM, and the fact patch coordinates are arbitrary (we could have rotated metapopulations 90, 180 or 270° without changing anything meaningful), it is better to calculate every metric of interest replicate by replicate first and only average later. It is important to note that the inclusion of temporal covariance matrices generates what are *de facto* observation-level random effects (Harrison 2014), so there is no further overdispersion to account for.

The intuition behind this formulation (latent residuals via OLRE + specified covariance structure) is the same as in this comment by Paul Bürkner on the brms R package development page (https://github.com/paul-buerkner/brms/issues/600#issuecomment-511677732).

The model is *de facto* a multivariate model, with a separate response for each $x, y$ combination. However, it can just as easily be rewritten as an univariate model for fitting purposes, without changing anything to the multivariate nature of the underlying model (in particular, the among-patches temporal correlations are preserved). The univariate re-formulation makes it easier for some packages to estimate some parameters (here the fixed effects ones); this formulation is the one we provide below.

The formula for the model for the number of adult females $N_{i,x,y,t}$ in metapopulation $i$, in the patch of coordinates $x, y$ at time $t$ is

$$N_{[i,x,y,t]} \sim \text{Poisson}(\lambda_{[i,x,y,t]}),$$

$$\log(\lambda_{[i,x,y,t]}) = B_0 + \sum_{j=1}^{J}(B_j \times x_{j[i,x,y]}) + a_{[i]} + c_{[i,x,y]} + d_{[i,x,y,t]},$$

$$a_{[i]} \sim \text{Normal}(0, \sigma_a),$$

$$c_{[i,x,y]} \sim \text{Normal}(0, \sigma_c),$$

$$\begin{bmatrix} d_{[i,1,1,t]} \\ ... \\ d_{[i,3,3,t]} \end{bmatrix} \sim \text{MVNormal}\left(\begin{bmatrix} 0 \\ ... \\ 0 \end{bmatrix}, \boldsymbol{\Omega}_{[i]}\right),$$

$$\boldsymbol{\Omega}_{[i]} = \begin{bmatrix} \sigma_{d[i,1,1]} & 0 & ... \\ 0 & \ddots & \\ \vdots & & \sigma_{d[i,3,3]} \end{bmatrix} \boldsymbol{R}_{[i]} \begin{bmatrix} \sigma_{d[i,1,1]} & 0 & ... \\ 0 & \ddots & \\ \vdots & & \sigma_{d[i,3,3]} \end{bmatrix},$$

where $B_j$ are the fixed effects (with $B_0$ the intercept), $a$ are replicate/metapopulation random effects, $c$ patch-level random effects, and $d$ temporal abundance fluctuations (We use Latin letters here, rather than the usual Greek script, to avoid confusion with $\alpha$, $\beta$, $\gamma$ variabilities). $\boldsymbol{\Omega}_{[i]}$ is the temporal covariance matrix for the replicate $i$ and $\boldsymbol{R}_{[i]}$ the corresponding correlation matrix. For implementation, we transform the treatment covariates into dummy centred variables following Schielzeth (2010), this has the added benefit of making $B_0$ the intercept of the "average" treatment.

All key metrics of interest in the present paper can be derived from this model:

- the mean local population size of the patch $x, y$ in metapopulation $i$ is $\mu_{i,x,y} = \exp(B_0 + \sum_{j=1}^{J}(B_j \times x_{j[i,x,y]}) + a_{[i]} + c_{[i,x,y]} + \frac{\sigma^2_{d[i,x,y]}}{2})$, i.e. the back-transformed sum of the patch-level latent mean and half the within-patch latent variance (based on the formula for the mean of the log-normal distribution; see also Villemereuil et al. 2016). Note that because the observed-scale mean depends on both the latent scale mean and the latent scale within-patch variance, and because our model allows both to vary between treatments, one cannot make inferences about treatment effects on means based on fixed effects alone;
- the observed-scale temporal variances and covariances $w$ (to use the syntax in main text **Table 1**) can be derived from the latent scale patch-level means $(B_0 + \sum_{j=1}^{J}(B_j \times x_{j[i,x,y]}) + a_{[i]} + c_{[i,x,y]})$ and latent scale within-patch variances $(\sigma^2_{d[i,x,y]})$ using the procedures detailed in Villemereuil et al. (2016);
- $\alpha$, $\beta_2$, $\varphi$ and $\gamma$ can be estimated for each metapopulation as outlined in main text **Table 1** once one has all the $\mu$ and $w$ corresponding to that metapopulation.

#### Meta-population model

Similarly, we can write a (much simpler) model for the total metapopulation size (total number of adult females counted in the metapopulation at one time step) $M$. Because there are here no internal spatial correlations to worry about here, we can use a negative binomial model here to model the within-replicate temporal variation:

$$M_{[i,t]} \sim \text{NegBinomial}(\lambda_{[i]}, \phi[i]),$$

$$\log(\lambda_{[i]}) = B_0 + \sum_{j=1}^{J}(B_j \times x_{j[i]}) + c_{[i]},$$

$$\log(1/\phi_{[i]}) = a_0 + \sum_{j=1}^{J}(a_j \times x_{j[i]}) + d_{[i]},$$

$$c_{[i]} \sim \text{Normal}(0, \sigma_c),$$

$$d_{[i]} \sim \text{Normal}(0, \sigma_d).$$

Here $c$ and $d$ refer to the metapopulation-level random effects, for the mean parameter and for the shape parameter, respectively. Similarly, $B$ and $a$ refer to the fixed effects coefficients for the mean and the shape. We fit the model for the overdispersion parameter on its log-transformed inverse: the inverse transformation (as suggested in the Stan language wiki: https://github.com/stan-dev/stan/wiki/Prior-Choice-Recommendations) is used to avoid giving too much prior weight to high overdispersion, the log transformation to keep $\phi$ estimates $> 0$.

#### Rationale behind priors

Given the models complexity, we combined general weakly informative priors sensu McElreath (2020) with more informative priors based on preexisting density data from a previous study (De Roissart et al. 2015, 2016), which we multiplied as needed to match the total area of a patch/a metapopulation. Ignoring these prior sources of information led to models predicting consistently too high abundances (not shown), although relative differences between treatments remained qualitatively similar.

In the patch-level model, priors for the fixed-effects coefficients $\beta_j$ (except $\beta_0$) and for the random effect correlation matrices followed McElreath's suggestions (here $\text{Normal}(0, 1)$ and $\text{LKJCorr}(2)$ respectively). Priors for the intercept $\beta_0$ and for the random effect standard deviations were based on the whole distribution of prior abundance data (De Roissart et al. 2015, 2016) and on the variance of that distribution, both on the log scale. We used a $\text{Normal}(2.8, 1)$ prior for $\beta_0$, and a $\text{Half}-\text{Normal}(0, 0.5)$ for the $\sigma$ parameters (when summing across the different random effect levels, this gives an overall prior for log-scale total variance centred on roughly 1, matching prior information).

In the metapopulation-level model, we replaced the prior for $\beta_0$ by $\text{Normal}(5, 1)$ in order to match the fact that metapopulations contain 9 patches, and used general purpose $\text{Normal}(0, 1)$ priors for all fixed-effects parameters linked to the shape $\phi$.

### S06 – Posterior pairwise comparisons

The following plots describe the outcome of posterior pairwise comparisons between treatments, regarding local population sizes (**Figs S06-1** to **S06-3**, match with main text **Figs 1-2**) and variability metrics (**Figs S06-4** and **S06-5**, match with main text **Fig. 4**). In all cases, comparisons are ratios, and dots and segments represent the posterior mean and 95% Highest Density Interval.

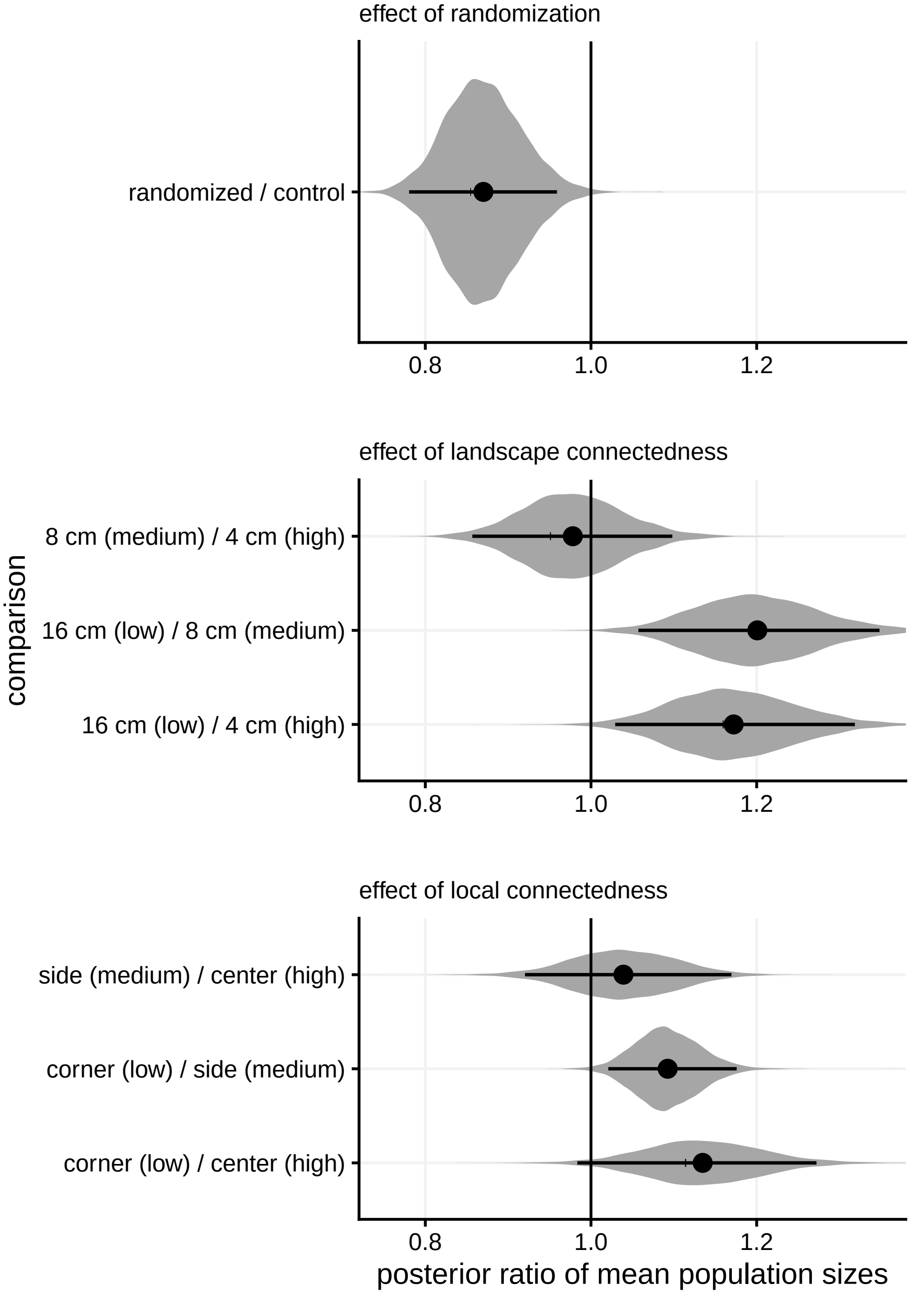


**Figure S06.1** – Additive effects of connectedness and randomization on local population size. For each treatment variable the effects of the others are pooled.

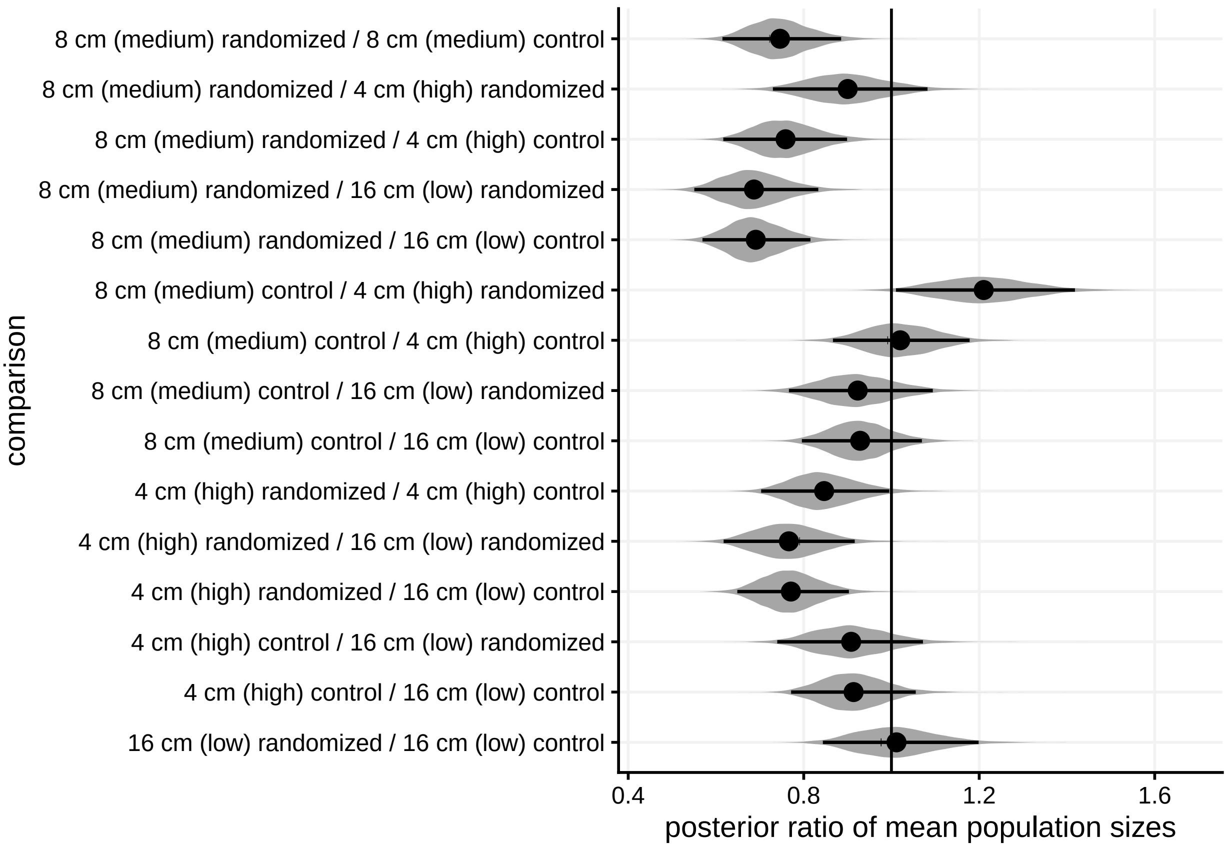


**Figure S06.2** – Combined effects of metapopulation-level connectedness and randomization on local population size.

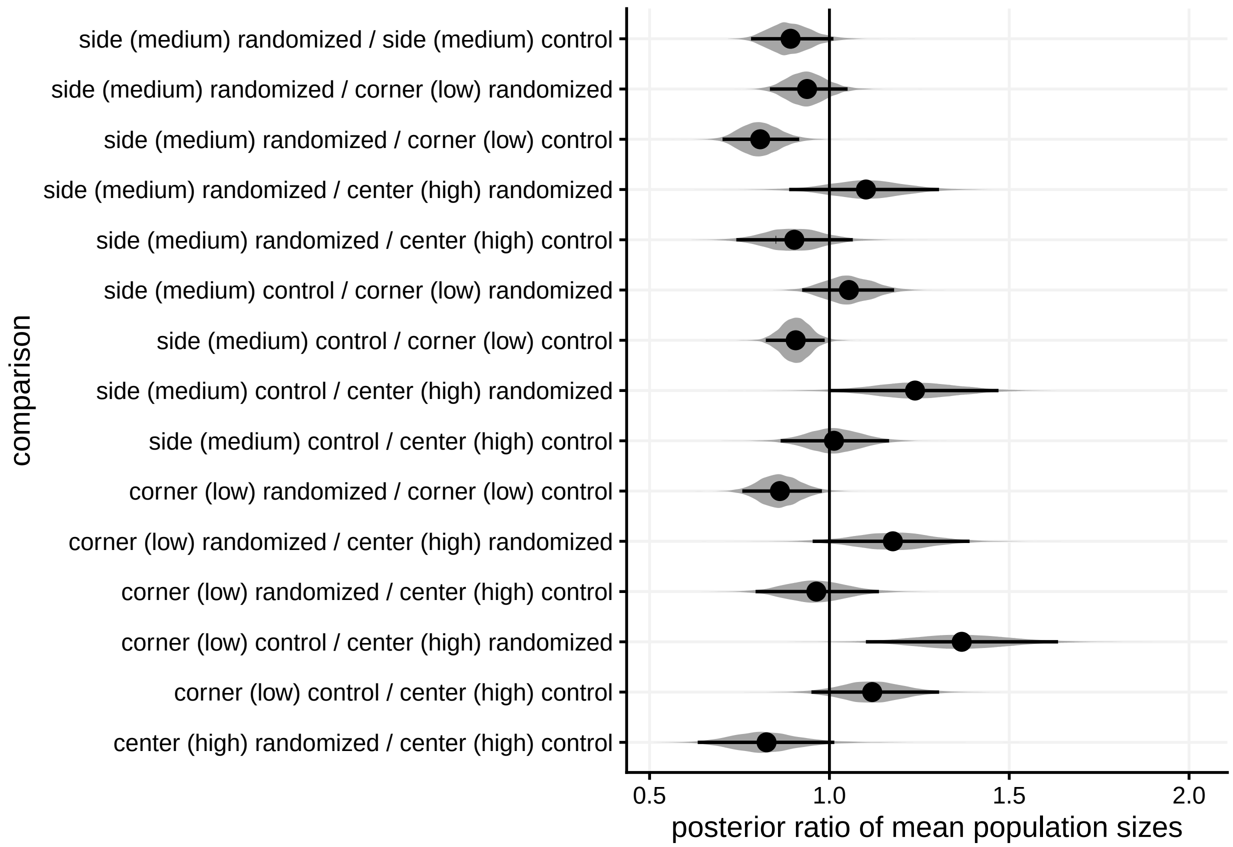


**Figure S06.3** – Combined effects of local connectedness and randomization on local population size.

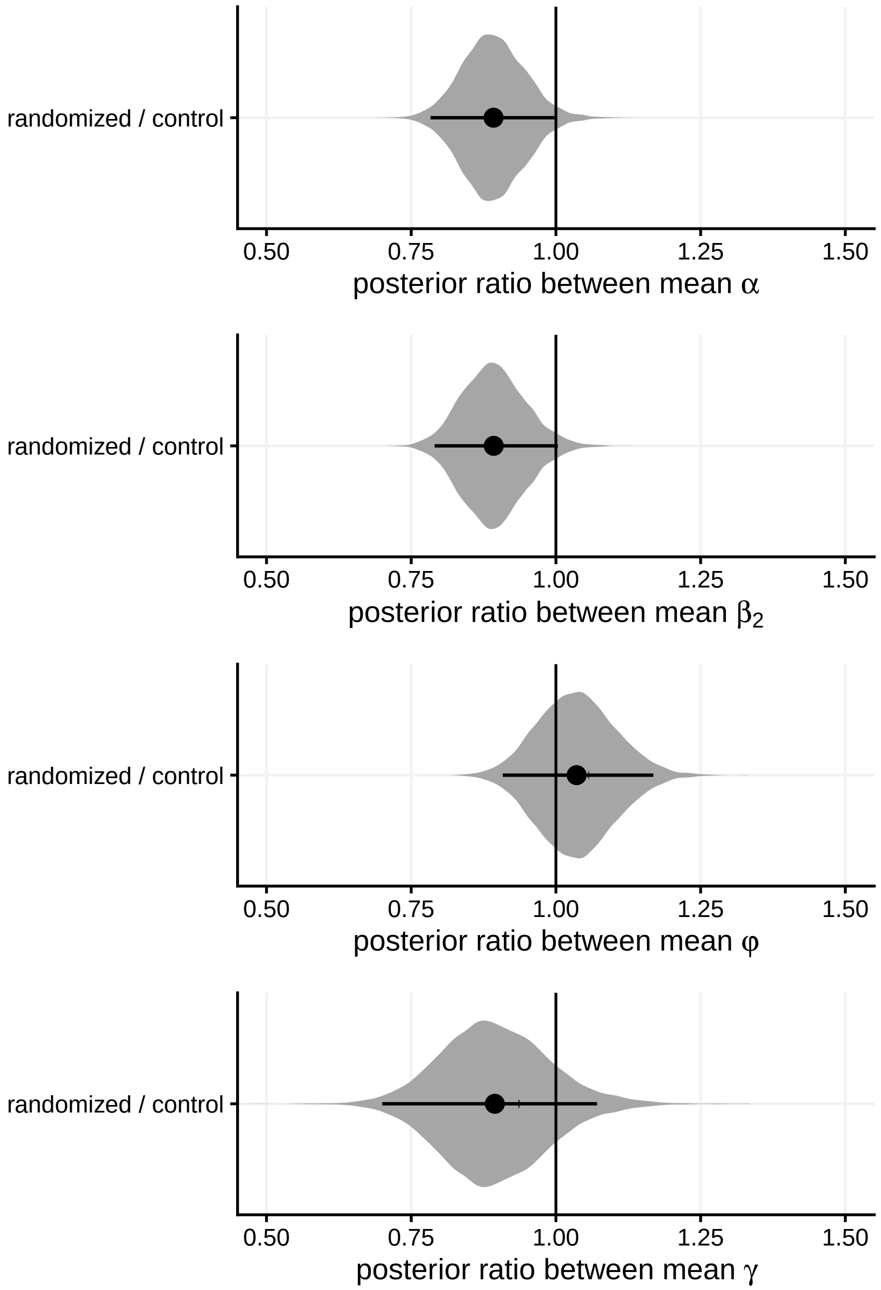


**Figure S06.4** – Effect of randomization on variability metrics

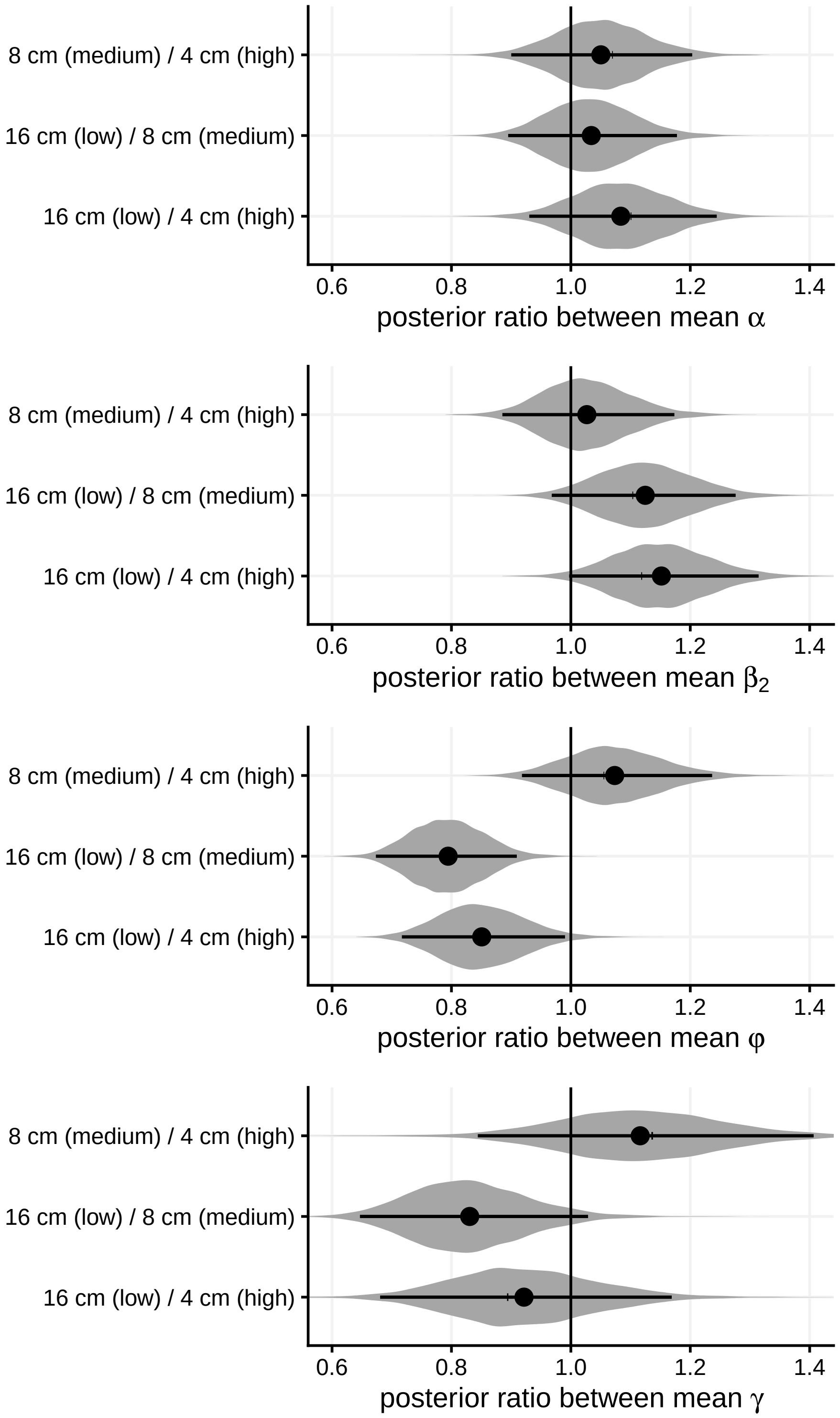


**Figure S06.5** – Effect of metapopulation-level connectedness on variability metrics

### S07 – Comparisons between patch-level model predictions, metapopulation-level model predictions, and observed data

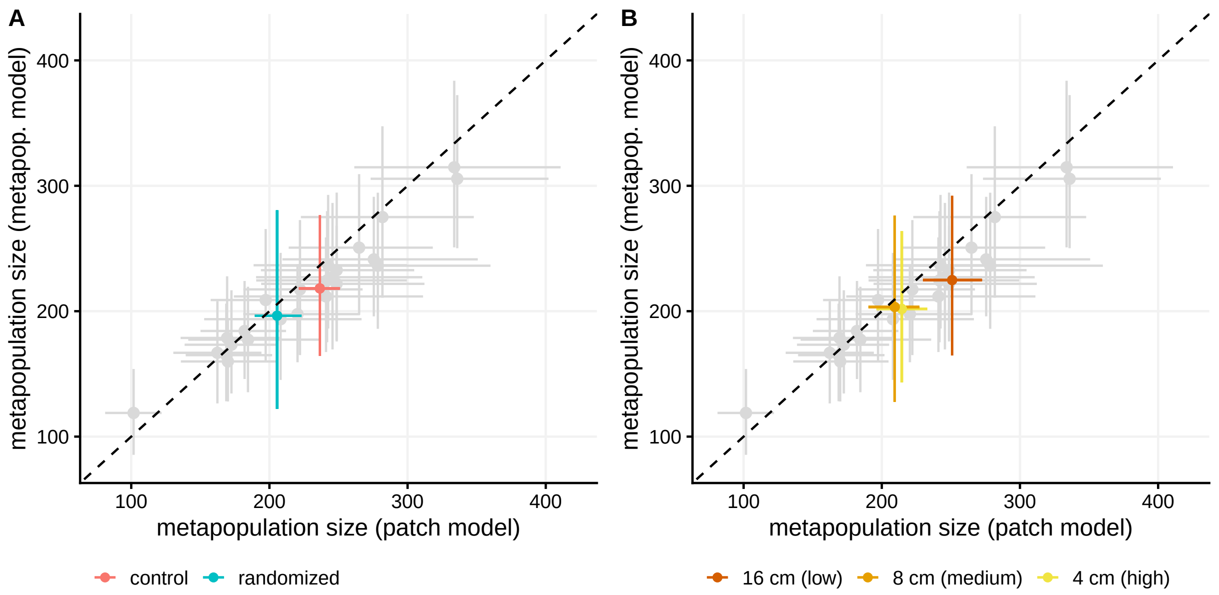


**Figure S07.1** – Comparisons between mean metapopulation size predictions (with 95% intervals) from the metapopulation-level model vs. the patch-level model (multiplying mean predictions by the number of patches, implicitly assuming patches are temporally independent). Treatment-level predictions (A: randomization treatment, B: metapopulation connectedness treatment) are added over replicate-level predictions (in grey). Dotted line: $y = x$.

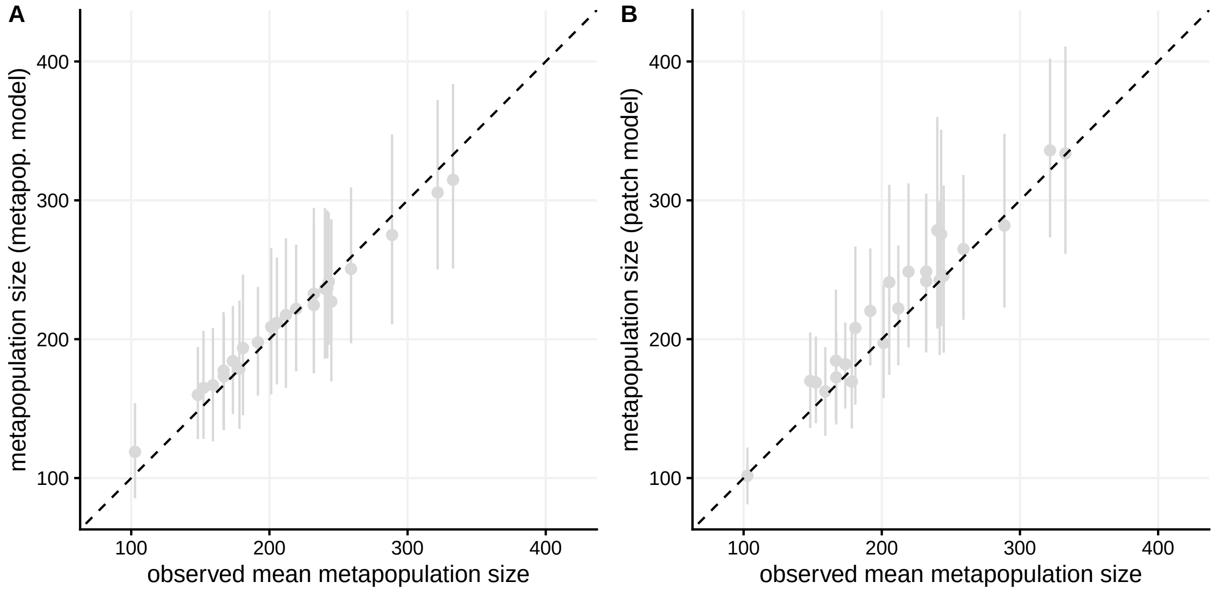


**Figure S07.2** – Comparisons between observed mean metapopulation sizes and mean metapopulation size predictions (with 95% intervals) from the metapopulation-level model (A) or the patch-level model (B: multiplying mean predictions by the number of patches, implicitly assuming patches are temporally independent). Dotted line: $y = x$.
